# Platelet Factor 4 (PF4) Regulates Hematopoietic Stem Cell Aging

**DOI:** 10.1101/2024.11.25.625252

**Authors:** Sen Zhang, Charles E. Ayemoba, Anna M. Di Staulo, Kenneth Joves, Chandani M. Patel, Eva Hin Wa Leung, Sang-Ging Ong, Claus Nerlov, Maria Maryanovich, Constantinos Chronis, Sandra Pinho

## Abstract

Hematopoietic stem cells (HSCs) responsible for blood cell production and their bone marrow regulatory niches undergo age-related changes, impacting immune responses and predisposing individuals to hematologic malignancies. Here, we show that the age-related alterations of the megakaryocytic niche and associated downregulation of Platelet Factor 4 (PF4) are pivotal mechanisms driving HSC aging. PF4-deficient mice display several phenotypes reminiscent of accelerated HSC aging, including lymphopenia, increased myeloid output, and DNA damage, mimicking physiologically aged HSCs. Remarkably, recombinant PF4 administration restored old HSCs to youthful functional phenotypes characterized by improved cell polarity, reduced DNA damage, enhanced *in vivo* reconstitution capacity, and balanced lineage output. Mechanistically, we identified LDLR and CXCR3 as the HSC receptors transmitting the PF4 signal, with double knockout mice showing exacerbated HSC aging phenotypes similar to PF4-deficient mice. Furthermore, human HSCs across various age groups also respond to the youthful PF4 signaling, highlighting its potential for rejuvenating aged hematopoietic systems. These findings pave the way for targeted therapies aimed at reversing age-related HSC decline with potential implications in the prevention or improvement of the course of age-related hematopoietic diseases.

**Key Points:**

- Age-related attrition of the megakaryocytic niche and associated PF4 downregulation is a central mechanism in HSC aging.
- PF4 supplementation, acting on LDLR and CXCR3 receptors, rejuvenates the function of aged HSCs.

## Introduction

Aging is generally considered an inevitable and irreversible process during the mammalian lifespan, concomitant with functional tissue deterioration and increased disease incidence^1^. In the blood system, hematopoietic stem cells (HSCs), responsible for lifelong blood cell production, also undergo age-related phenotypical and functional decline^2^. Specifically, HSC aging is characterized by the expansion of less-functional HSCs biased towards myeloid blood cell lineages, deregulated self-renewal capacity, and impaired regenerative potential. These aging-associated alterations lead to compromised immune responses, frequently accompanied by clonal hematopoiesis, potentially underlying leukemic transformation^3,4^. Notably, the aging of the human hematopoietic system mirrors many of the phenotypes observed in murine models. HSCs from old individuals display increased cell frequencies, reduced lymphoid output, and a higher prevalence of platelet-biased HSCs with transcriptional priming toward platelet lineage differentiation compared to young HSCs^5–7^. Mechanistically, HSC aging has been associated with cell-intrinsic processes such as DNA damage accumulation, epigenetic modifications, mitochondrial DNA mutations, and metabolic dysfunction^8^. Nevertheless, it is now widely recognized that the bone marrow HSC regulatory niche or microenvironment also undergoes extensive age-related alterations that contribute to, or even drive, the aging phenotype of HSCs^9,10^.

In homeostasis, HSCs reside within specialized bone marrow niches that regulate their self-renewal, quiescence, localization, differentiation, and overall function, ensuring blood cell production while preventing HSC exhaustion^2^. However, the nature and extent of age-related alterations that each bone marrow niche undergoes and how these changes contribute to specific features of HSC aging remain poorly understood. The bone marrow niche comprises various cell types, including stromal niche-supporting cells like endothelial and mesenchymal stem cells (MSCs) and feedback signals from HSC progeny that regulate HSC activity. Among these niche constituents, megakaryocytes (MKs) play a crucial role in maintaining HSC quiescence via the secretion of platelet factor 4 (PF4, also known as CXCL4)^11^ and transforming growth factor-β (TGF-β)^12^. Studies in young mice further revealed that the localization, proliferation, and reconstitution potential of platelet- and myeloid-biased HSCs, the HSC subset that selectively expands with age^13,14^, is regulated by MKs, while periarteriolar MSC niches mainly regulate balanced/lymphoid-biased HSCs^15^. Recent studies have revealed alterations in the aged MK niche, where MK progenitor (MkP) cell function is enhanced upon aging^16^, and aged HSCs are redistributed away from old MK niches^17,18^, the opposite of young HSCs^11,12^. Similarly, human hematopoietic stem and progenitors show reduced interactions with MKs in human bone marrow, suggesting conserved aging-associated niche phenotypes across species^19^. Still, the mechanisms underlying the loss of association between aged HSCs and MKs in the aged bone marrow remain unclear. Whether changes in the aged megakaryocytic niche and its secreted factors directly contribute to HSC aging remains an open question. Here, we identified the attrition of the megakaryocytic niche and the concomitant decline in PF4 levels as key mechanisms driving HSC aging. We also explored the potential of PF4 as a factor for HSC rejuvenation and anti-aging interventions in the hematopoietic system.

## Methods

### Mice

Young (2-3 months) and aged (18-22 months) C57BL/6-CD45.1 (B6.SJL-Ptprc^a^ Pepc^b^/BoyJ) and C57BL/6-CD45.2 congenic strains were purchased from Jackson Laboratory or by aging young mice in our facilities. B6.129S7-Ldlr^tm1Her^/J (*Ldlr^−/−^*)^20^, C57BL/6-Tg(Pf4-iCre)Q3Rsko/J (*Pf4-Cre*)^21^, C57BL/6-Gt(ROSA)26Sor^tm1(HBEGF)Awai^/J (iDTR)^22^, B6.129P2-Cxcr3^tm1Dgen^/J (*Cxcr3^−/−^)* (strain#005796) were generated by Deltagen and purchased from Jackson Laboratory. *Pf4^−/−^* mice^23^ and *vWF-eGFP* transgenic mice^24^ were bred in our facilities. B6.129S7-Ldlr^tm1Her^/J and B6.129P2-Cxcr3^tm1Dgen^/J mice were crossed to generate B6.129S7-Ldlr^tm1Her^/J; B6.129P2-Cxcr3^tm1Dgen^/J mice (DKO mice) in our facilities. Sex-matched animals from the same age group were randomly assigned to all experimental groups. This study complied with all ethical regulations involving experiments with mice, and all experimental procedures were approved by the Office of Animal Care and Institutional Biosafety (OACIB) at the University of Illinois at Chicago.

### Immunofluorescence staining and imaging of sorted HSCs

Immunofluorescence staining for γH2AX or cdc42 and α-tubulin was performed as described^18^. All images were acquired using a Zeiss LSM880 Confocal Microscope (Zeiss) and reconstructed in 3D or 2.5D with ZEN software (Zeiss). Image analysis was performed using ImageJ2 2.14.0/1.54f.

### Gene expression analysis of young and old MK

Differential gene expression analysis was performed to identify age-associated MK genes. The generated datasets were deposited in GEO and are available under GSE280042.

Additional Materials and Methods are provided in **Supplemental Methods.**

## Results

### Aging induces remodeling and impairment of the megakaryocytic niche function

To investigate the age-associated alterations of the megakaryocytic niche, we first analyzed the bone marrow of young (2-3 months old) and old (20-24 months old) von Willebrand factor (vWF)-eGFP mice^24^, by whole-mount sternum 3D imaging. The number of MK-lineage cells identified by their size, morphology, and CD150^+^ vWF^+^ increases with age (**Figure 1A-1B**). FACS analyses confirmed our imaging results and revealed the increased frequency and number of FSC^high^ CD41^+^CD150^+^ MKs (**Figure 1C-1D**) and Lineage (Lin)^-^ Sca-1^-^c-Kit^+^CD150^+^CD41^+^ MK progenitors (MkP, **Figures 1E-1F**), as reported^16–18^. Consistent with a recent study describing morphologic changes in human MKs with age^19^, old murine MK-lineage cells are significantly smaller than their young counterparts (∼23±5.2 μm *vs* ∼26±6.8 μm diameter, respectively), suggesting potential alterations in their maturation status (**Figure 1G**), which is associated with MK niche function^25^. Flow cytometry ploidy analyses further revealed a significant decrease in the percentage of polyploid MKs (>4N) in the bone marrow of old mice (**Figures 1H-1I**).

**Figure 1.**
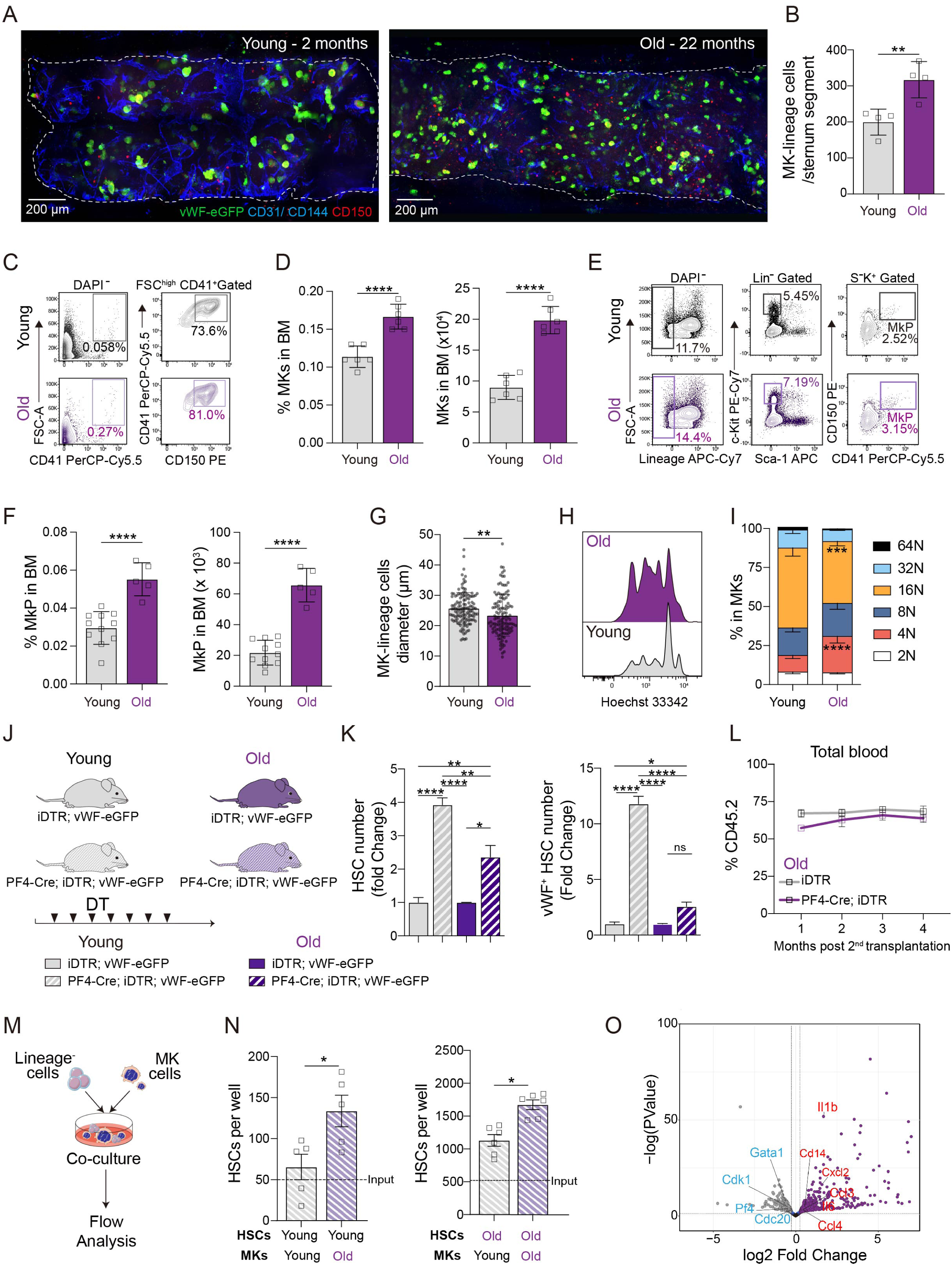
Aging induces remodeling of the megakaryocytic niche. **A.** Whole-mount confocal images of the sternal bone marrow (BM) from young (2 months old) and old (22 months old) *vWF-eGFP* mice. MK-lineage cells are distinguished by their size, morphology, CD150, and vWF expression. Endothelial cells are labeled with CD31/CD144. **B.** Quantification of MK-lineage cell (CD150^+^ vWF^+^) number from the sternal BM of young (2-3 months) and old (20-24 months) mice (young n = 4, old n = 4). **C.** Representative flow cytometry analysis of MK (FSC^high^ CD150^+^ CD41^+^) in the BM of young and old mice. **D.** Percent (left) and absolute number (right) of MKs found in the BM of young and old mice (young n = 6, old n = 6). **E.** Representative flow cytometry analysis of MK progenitors (MkP, Lin^-^ c-Kit^+^ Sca-1^-^ CD150^+^ CD41^+^) in the BM of young and old mice. **F.** Percent (left) and absolute number (right) of MkPs found in BM of young and old mice (young n = 11, old n = 5). **G.** Average MK-lineage cell diameter from the sternal BM of young and old mice. **H.** Representative histogram of young and old MK ploidy distribution and **I.** quantification (young n = 5, old n = 6). **J.** Experimental design to test the effect of MK depletion on young and old mice. **K.** Fold change of HSC and vWF^+^ HSC numbers after MK depletion in young and old mice. **L.** Total blood chimerism (CD45.2^+^) of secondary recipient mice transplanted with BM cells from 1^st^ recipients, which received HSCs from old *iDTR* and old *PF4-Cre; iDTR* mice after 7-day diphtheria toxin (DT) injection. **M.** Experimental outline of MKs and HSCs co-culture. CD45.2^+^ young or old Lineage^-^ cells were co-cultured with CD45.1^+^ young or old MKs for 4 days. **N.** Flow cytometric analysis of CD45.2^+^ Lin^-^ c-Kit^+^ Sca-1^+^ CD150^+^ CD48^-^ HSC numbers obtained under various conditions. The starting number of HSCs seeded in the co-culture is indicated (Input). **O.** Volcano plots indicating differentially expressed genes (DEGs, P < 0.05, Fold change > 1.2) between young and old MKs. Data are presented as mean ± SD. *P < 0.05, **P < 0.01, ***P < 0.001, ****P < 0.0001.

To assess the role of old MKs in HSC regulation *in vivo*, we depleted MKs in young and old *Pf4-Cre*;*iDTR*;*Vwf-eGFP* and control *iDTR*;*Vwf-eGFP* mice with diphtheria toxin (DT) (**Figure 1J**), as described^11,15^. Our previous results in young mice revealed that MK depletion leads to a drastic increase in the number of phenotypic and functional HSCs, in particular, the platelet/myeloid-biased HSC subset marked by vWF expression^11,15^. As expected, MK depletion significantly increased the total number of bone marrow Lin^-^ Sca-1^+^ c-Kit^+^ CD135^-^ CD48^-^ CD150^+^ HSCs (by 3.92-fold) in young mice, especially the myeloid-biased vWF^+^ HSC subset (by 11.79-fold; **Figure 1K**). In contrast, we found that in old mice, MK depletion only leads to a modest expansion of total and vWF^+^ HSCs (2.08-fold and 2.39-fold, respectively) (**Figure 1K**). In agreement with these results, primary and secondary competitive transplantation assays of sorted HSCs from old *Pf4-Cre*;*iDTR* and control *iDTR* mice, 7 days after DT treatment, did not reveal any significant differences in donor reconstitution or tri- lineage differentiation (**Figure 1L; supplemental Figures 1A-1B**). These results suggest that aging impairs the niche quiescence-enforcing capacity of old MKs or that aged HSCs lost the capacity to respond to MK signals. To further investigate the functional changes in the HSC niche capacity of old MKs, we directly co-cultured young and old Lineage^−^ cells with young and old MKs (**Figure 1M**). Notably, in contrast to young MKs, old MKs fail to restrict the *in vitro* expansion of both young and old phenotypic HSCs, with a more pronounced effect in young HSCs (**Figure 1N**). Altogether, these results confirm that age-related alterations of the megakaryocytic niche led to an impairment of their HSC quiescence-enforcing capacity.

### Aging leads to deregulated signaling in aged MKs

To understand the molecular changes driving MK age-related alterations, we performed RNA-seq of MKs sorted from the bone marrow of young and old mice (**supplemental Figure 1C; supplemental Table 1**), as described^26^. We observed that many genes were differentially regulated by age in MKs, with 527 upregulated and 728 downregulated in old MKs (**Figure 1O**). Gene ontology analysis^27^ revealed that various immune and inflammatory process networks were enriched in old MK, portrayed by the upregulation of genes such as *IL-1b, Cd14, Ccl3, Ccl4, IL-6,* and *Cxcl2* (**Figure 1O; supplemental Figure 1D**; **supplemental Table 1**). These results suggest that age-related attrition of the megakaryocytic niche also contributes to the heightened and detrimental inflammatory microenvironment associated with physiological HSC aging^17,28–31^. Our analysis further revealed the age-associated downregulation of multiple genes associated with cell cycle progression, such as *Cdk1* and *Cdc20* and DNA synthesis (**Figure 1O; supplemental Figure 1D; supplemental Table 1),** two critical processes in MK terminal maturation and polyploidization^32^. In agreement with the more immature nature of old MK, the expression of *Gata1*, a transcriptional factor essential for normal megakaryopoiesis^33^, and *Pf4,* a known promoter of MK maturation via autocrine signaling^34,35^, was significantly downregulated in old mice (**Figure 1O, supplemental Figure 1D; supplemental Table 1)**. Real-time PCR (qPCR) experiments confirmed these results (**supplemental Figures 1E-1F**). Accordingly, PF4 protein levels were significantly downregulated in the serum of old mice (**supplemental Figure 1G**). In agreement, a recent study revealed that the plasma levels of PF4 decline in aged mice and in humans^36^. Given that PF4 is a critical factor for maintaining HSC quiescence and specifically regulates the myeloid/platelet-biased HSC compartment^15^ and whose expression is almost exclusively restricted to MKs and MK-derived platelets^21^, these results suggest that PF4 may be a key determinant in HSC aging.

### Young *Pf4^−/−^* mice exhibit phenotypes reminiscent of premature HSC aging

To specifically evaluate the role of PF4 in hematopoietic aging, we assessed HSC aging hallmarks in PF4-deficient (*Pf4^−/−^*) mice^23^. Our previous studies revealed increased HSC number and loss of HSC−MK association in young *Pf4^−/−^* mice^11^, two phenotypes emulating physiological HSC aging. Outside of the hematopoietic system, a recent study further revealed that middle-aged *Pf4^−/−^* mice demonstrate accelerated cognitive aging^36^. Our analyses revealed increased myeloid and decreased lymphoid (B and T cells) output in the bone marrow of young *Pf4^−/−^* mice (2-3 months), as seen in an aged immune system (**Figure 2A**-**2B**). We also found increased platelet numbers in the blood, as reported ^23^, mirrored by an increase in bone marrow MkPs (**supplemental Figures 2A-2B**). However, the number of bone marrow MK, peripheral white blood cells (WBC), and red blood cells (RBC) in *Pf4^−/−^*mice was unchanged (**supplemental Figures 2C-2E**). As seen in old MK (**Figure 1N**), young *Pf4^−/−^*MK also display an impaired capacity to restrict the *in vitro* expansion of young HSCs in direct co-culture conditions (**Figure 2C**). Notably, young *Pf4^−/−^* HSCs also show elevated levels of DNA damage (identified by γH2AX foci; **Figure 2D**) and loss of polar distribution of α- tubulin and cdc42 (**Figures 2E**-**2F, supplemental Figures 2F-2G)**, which are strongly linked to the loss of HSC function^37^ and lymphoid potential^38,39^ in old HSCs^40,41^.

**Figure 2.**
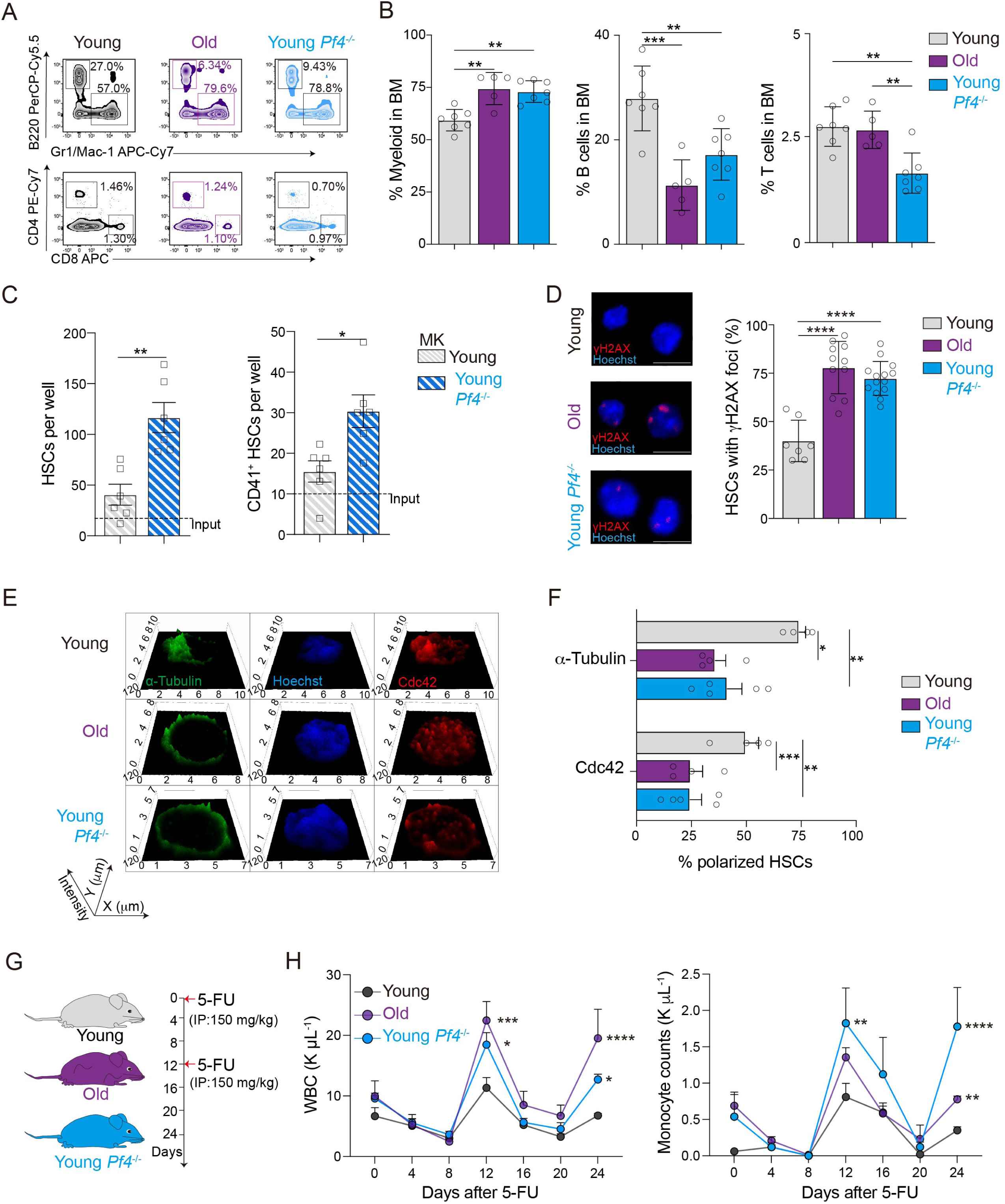
Young *Pf4^-/-^*mice exhibit phenotypes reminiscent of premature HSC aging. **A.** Representative FACS plots of myeloid, B, CD4^+^, and CD8^+^ T cells in the BM of young (2-month-old), old (22-month-old), and young *Pf4^-/-^* (2-month-old) mice. **B.** Percentage of myeloid, B, and T cells (combined CD4^+^ and CD8^+^ T cells) in the BM of young, old, and *Pf4^-/-^* mice (young n = 7, old n = 5, young *Pf4^-/-^* n = 7). **C.** CD45.2^+^ young Lineage^-^ cells were co-cultured with CD45.1^+^ MK cells from young control and young *Pf4^-/-^*mice for 4 days. Flow cytometric analysis of CD45.2^+^ total HSCs and CD41^+^ HSCs. The starting number of HSCs seeded in the co-culture is indicated (Input). **D.** Left: Representative confocal z-stack projections of HSCs sorted from young, old, or young *Pf4^-/-^* mice stained for γH2AX and Hoechst. Scale bar, 10 µm. Right: Quantification of the percentage of HSCs with γH2AX foci (calculated as the mean percentage of a total of 181, 264, and 607 HSCs isolated from 3 young, 3 old, and 4 young *Pf4^-/-^* mice). **E.** Representative confocal 2.5 D images of HSCs sorted from young, old, or young *Pf4^-/-^* mice and stained against cdc42, α-Tubulin, and Hoechst. **F.** Quantification of the percentage of cdc42 and α-Tubulin polarized HSCs out of total HSCs scored (total of 53, 60, 64 HSCs isolated from 3 young, 3 old, and 4 young *Pf4^-/-^* mice). **G.** Experimental design of 5-FU injection (IP, Intraperitoneal injection). **H.** Regeneration of peripheral white blood cells (WBC) and monocytes after 5-FU treatment of young, old, or young *Pf4^-/-^* mice (young n = 3, old n = 3, young *Pf4^-/^*^-^ n = 6). Data are mean ± SEM. *P < 0.05, **P < 0.01, ***P < 0.001, ****P < 0.0001.

While *Pf4^−/−^* mice are phenotypically normal and live normal lifespans under homeostatic laboratory conditions^23^, how these mice cope with stress is unknown. We next investigated *Pf4^−/−^* mice’s response to myeloablative stress. Young, old, and young *Pf4^−/−^*mice were injected with 5-fluorouracil (5-FU; **Figure 2G**). Old mice are known to be highly sensitive to the 5-FU challenge^30,42^. Time course blood analyses revealed that in contrast to young control mice, young *Pf4^−/−^*mice mirror the exacerbated production of WBCs, including monocytes, neutrophils, and lymphocytes, similar to the response seen in old mice after 5-FU (**Figure 2H**; **supplemental Figure 2H**). Furthermore, young *Pf4^−/−^*mice are more sensitive to 5-FU, with increased mortality compared to control young mice (**supplemental Figure 2I),** suggesting impaired hematopoietic recovery. Accordingly, PF4 has been shown to be important in limiting the duration of the regenerative process during emergency myelopoiesis^43^. These results indicate that loss of PF4 accelerates hematopoietic aging and that PF4 might be a key factor in HSC rejuvenation strategies.

### PF4 administration reverses HSC aging phenotypes

Since our results indicate that PF4 levels are critical in HSC aging, we next investigated the rejuvenation potential of PF4. We first tested if old HSCs retained the capacity to respond to PF4 using *in vitro* culture conditions. Interestingly, short-term recombinant PF4 treatment of old HSCs in culture decreased the percentage of γH2AX positive cells (20% reduction; **supplemental Figure 3A**). After a 4-day culture with PF4, the proliferation of old HSCs, including old myeloid-biased CD41^+^ HSCs, was significantly inhibited, as seen by absolute cell counts and cell cycle analyses (**Figures 3A-3C**). These results indicate that old HSCs retain the potential to respond to PF4, as seen in young HSCs^11,15^.

**Figure 3.**
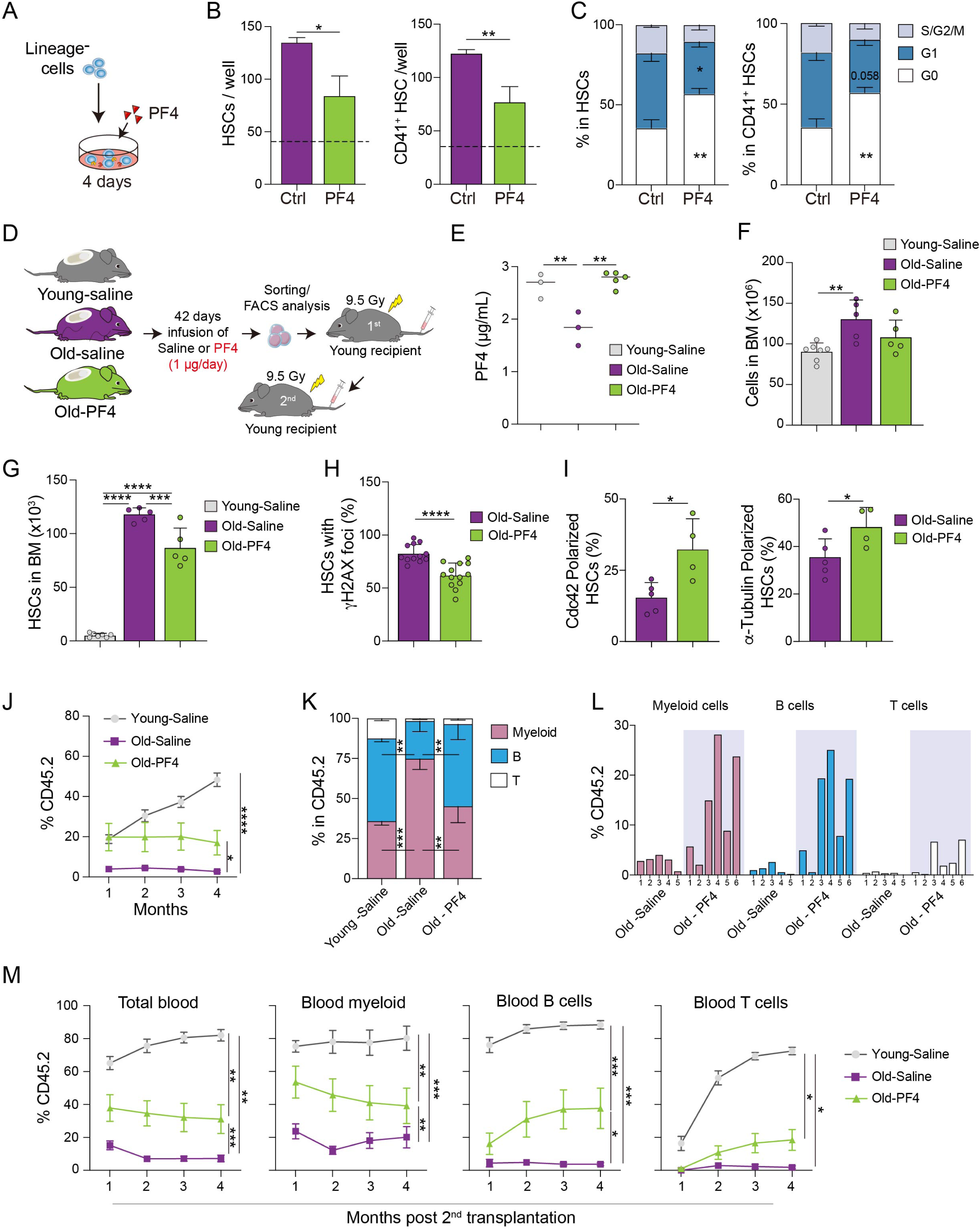
PF4 administration reverses HSC aging phenotypes. **A.** Outline of *in vitro* experimental strategy. **B.** Number of total (left graph) and myeloid-biased (CD41^+^ HSCs, right graph) old HSCs per well after culture with PF4 (young n = 6, old n = 6). **C.** Ki67 cell cycle analysis of total and myeloid-biased old HSCs cultured with PF4 (young n = 6, old n = 6). **D.** *In vivo* PF4 administration experimental design. **E.** PF4 levels in the serum of young-saline, old-saline, and old-PF4 mice (Young-Saline n = 3, Old-Saline n = 3, Old-PF4 mice n = 5, each dot is the mean of two replicates). **F.** Total cell counts in the BM of young-saline, old-saline, and old-PF4 mice (Young-Saline n = 7, Old-Saline n = 5, Old-PF4 mice n = 5). **G.** Total HSC number in the BM of young-saline, old-saline, and old-PF4 mice (Young-Saline n = 7, Old-Saline n = 5, Old-PF4 mice n = 5). **H.** Quantification of the percentage of HSCs with γH2AX foci (calculated as the mean percentage of a total of 387 and 185 HSCs isolated from 5 Old-Saline and 5 Old-PF4 mice). **I.** Quantification of the percentage of cdc42 (left graph) and α-Tubulin (right graph) polarized HSCs out of total HSCs scored (total of 138 and 103 HSCs isolated from 5 Old-Saline and 5 Old- PF4 mice). **J.** Total blood chimerism (CD45.2^+^) of CD45.1 recipient mice transplanted with 250 sorted HSCs from Young-Saline, Old-Saline, and Old-PF4 mice. **K.** Composition of myeloid, B, and T cells in the BM of CD45.2^+^ donor cells, 4 months after transplantation. **L.** Myeloid, B, and T cells chimerism (CD45.2^+^) in each individual CD45.1 recipient mice, 4 months after transplantation. **M.** Quantification of engraftment and tri-lineage (myeloid, B-cell, and T-cell) differentiation in the mice transplanted with 10 million total BM from 1^st^ recipients. Data are mean ± SEM. *P < 0.05, **P < 0.01, ***P < 0.001, ****P < 0.0001.

We thus tested the *in vivo* effects of recombinant PF4 administration on 22-month-old mice. Continuous systemic delivery of PF4 was achieved by the implantation of subcutaneous Alzet osmotic pumps for 42 days (**Figure 3D**), as described^18^. As controls, old and young mice also received pumps filled with saline. This regimen restored the serum levels of PF4 in old mice to those of young mice (**Figure 3E**). PF4 treatment reduced the absolute number of phenotypic HSCs in the bone marrow of old-PF4- treated mice (**Figures 3F**-**3G**), with no significant alterations in the number of other hematopoietic progenitor cell populations (data not shown). Importantly, PF4 treatment also reduced the levels of DNA damage and restored HSC polarity in old HSCs, compared to old-saline HSCs (**Figures 3H-3I).** Peripheral blood analyses further revealed an increase in the number of RBCs in old PF4-treated mice to levels comparable to young mice, while WBCs and platelet levels remained unchanged (**supplemental Figure 3B**). To gain further molecular insights into the HSC rejuvenation potential of PF4, we evaluated the expression levels of specific hematopoietic lineage genes previously identified as differentially expressed in old HSCs^44,18^. HSCs from old PF4-treated mice exhibited decreased expression of myeloid (*Fgr, Osmr,* and *Rab27b;* **supplemental Figure 3C**) and megakaryocyte/platelet (*Clu* and *Itgb3;* **supplemental Figure 3D**) lineage genes and increased expression of lymphoid (*Mtcp1* and *Blnk;* **supplemental Figure 3E**) lineage genes, mirroring the profile observed in young and balanced HSCs. Our findings suggest that PF4 imparts rejuvenated phenotypic and functional characteristics to old HSCs.

HSC aging is associated with myeloid-biased differentiation and decreased HSC reconstitution potential and long-term function^45^. To evaluate the long-term trilineage repopulation capacity of PF4-treated old HSCs, we performed *in vivo* primary and secondary competitive transplantations of sorted HSCs (**Figures 3J-3M**; **supplemental Figure 3F**). Notably, treatment with PF4 significantly increased old HSCs donor chimerism and improved lymphoid output 4 months following transplantation (**Figure 3J-3L**). Notably, the functional rejuvenation of old HSCs by PF4 treatment was also evident following secondary bone marrow transplantation, in which the enhanced engraftment and multilineage contribution persevered (**Figure 3M; supplemental Figure 3F**). Altogether, these data suggest that PF4 administration has the potential to attenuate and partially reverse HSC aging phenotypes and restore their functional properties.

### PF4 regulates HSCs through both CXCR3 and LDLR receptors

Previous studies showed that PF4 acts directly on HSCs to restrict their proliferation^11,15^, but the specific PF4 receptor(s) that promotes HSC quiescence remains unclear. LRP1 (low-density lipoprotein (LDL) receptor-related protein 1)^34^, EPCR (endothelial protein C receptor)^46^, CXCR2 (C-X-C motif chemokine receptor 2)^47,48^, LDLR (LDL receptor)^49^, CCR1 (C-C motif chemokine receptor 1)^50^, and CXCR3 (C-X-C Motif Chemokine Receptor 3)^36,51^ have been shown to transduce PF4 signals in a variety of cell types. Interestingly, these six candidate receptors are all expressed on both young and old HSCs at varying levels^18^ (**supplemental Figure 4A**). To investigate which PF4 candidate receptor was being used by HSCs, we first cultured Lineage^−^ cells in the presence or absence of PF4 and tested the effect of blocking antibodies against each receptor or the effect of an inhibitor specific to CCR1 (**Figure 4A**). Notably, only the blockade of LDLR and CXCR3 abrogated the anti-proliferative effect of PF4 on CD41^+^ HSCs *in vitro* (**Figure 4B**). To functionally validate the role of PF4-CXCR3 and PF4-LDLR in HSC proliferation, we used *Ldlr^−/−^* ^20^ and *Cxcr3^−/−^* ^52^ mice. Our results revealed that *Ldlr^−/−^* and *Cxcr3^−/−^* HSCs *in vitro* proliferation is not affected by PF4 treatment, as seen in control WT HSCs (**Figures 4C-4D**). These data confirm our blocking antibody experiments, suggesting that LDLR and CXCR3 are the PF4 receptors on HSCs regulating HSC proliferation.

**Figure 4.**
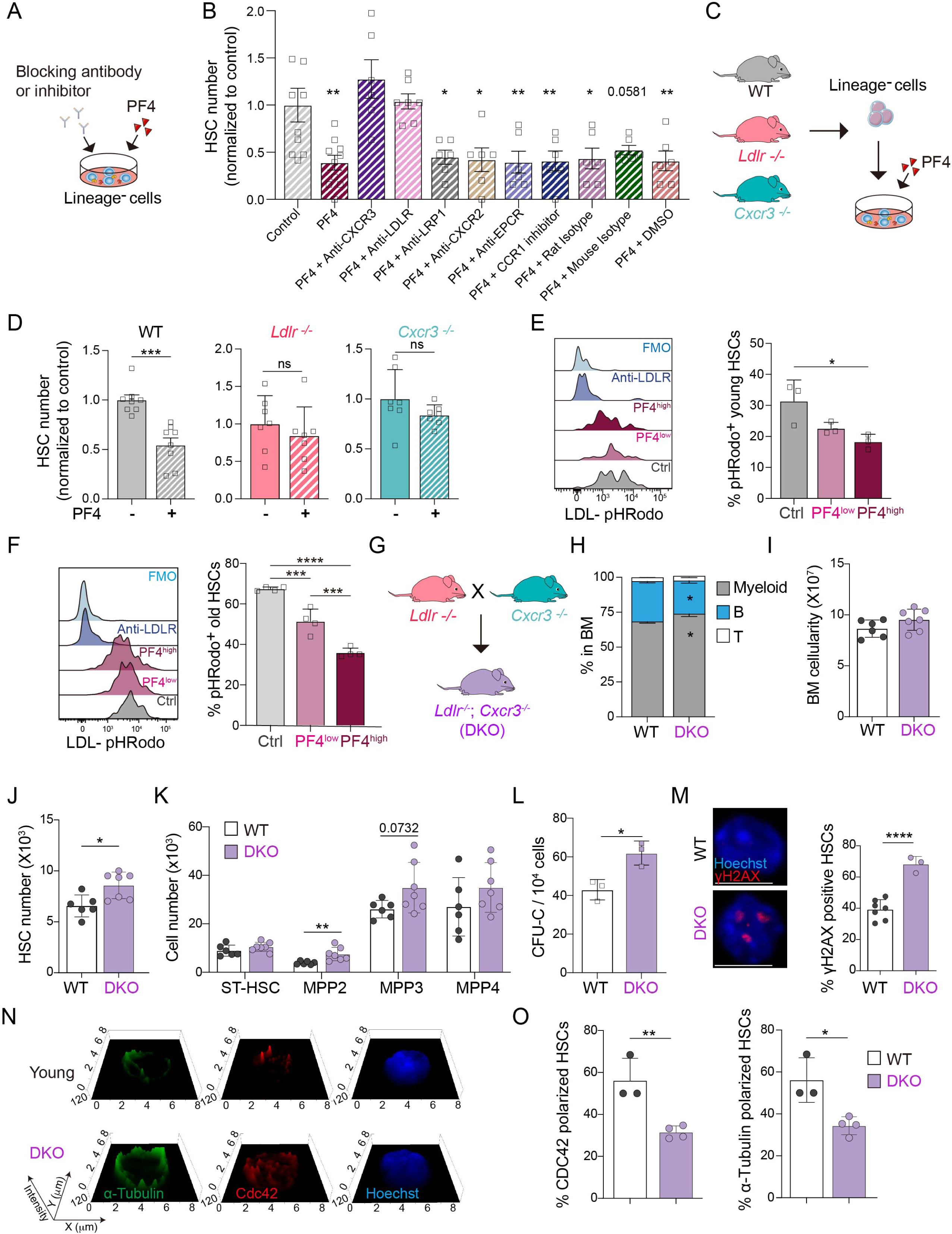
PF4 Regulates HSCs through both CXCR3 and LDLR receptors. **A.** Experimental strategy to investigate the PF4 receptor on HSCs. **B.** Normalized number of CD41^+^ HSCs to control well after culture with PF4 in the presence or absence of anti-LDLR, anti-CXCR3, anti-EPCR, anti-CXCR2 neutralizing antibodies, CCR1 inhibitor, or relevant controls (isotypes or DMSO). **C.** Experimental strategy to test the effect of PF4 on HSCs from *Ldlr^-/-^*and *Cxcr3^-^*^/-^ mice. **D.** Number of HSCs per well, normalized to control, after culture with PF4. **E.** Representative histogram and frequency of LDL-pHRodo+ young HSCs after PF4 treatment (PF4^low^: 1 μg/mL, PF4^high^: 5 μg/mL). Functional blocking anti-LDLR antibody was used as a negative control. **F.** Representative histogram and frequency of LDL-pHRodo+ old HSCs after PF4 treatment (PF4^low^: 1 μg/mL, PF4^high^: 5 μg/mL). **G.** Experimental strategy to generate DKO (*Ldlr^-/-^*; *Cxcr3^-/-^*) mice. **H.** Composition of myeloid, B, and T cells in the BM of young WT and DKO mice (WT n = 4, DKO n = 4). **I.** Total cell counts in the BM of young WT and DKO mice (WT n = 4, DKO n = 4). **J.** Absolute number of HSCs in BM of young WT and DKO mice (WT n = 6, DKO n = 7). **K.** Number of ST-HSC, MPP2, MPP3, and MPP4 in the BM of young WT and DKO mice (WT n = 6, DKO n = 7). **L.** Frequency of colony-forming unit cell (CFU-C) derived from 10^4^ BM cells from young WT and DKO mice. **M.** Left: Representative confocal z-stack projections of HSCs sorted from young WT and DKO mice and stained with γH2AX and Hoechst. Scale bar, 5 µm. Right: Quantification of the percentage of HSCs with γH2AX foci (calculated as the mean percentage of a total of 113 and 76 HSCs isolated from 3 young WT and 3 young DKO mice). **N.** Representative confocal 2.5 D images of HSCs sorted from young WT and DKO mice and stained with cdc42, α-Tubulin, and Hoechst. **O.** Quantification of the percentage of cdc42 and α-Tubulin polarized HSCs out of total HSCs scored (total of 49 and 72 HSCs isolated from 3 young WT and 3 young DKO mice). Data are mean ± SEM. *P < 0.05, **P < 0.01, ***P < 0.001, ****P < 0.0001.

LDLR is involved in the endocytosis of LDL cholesterol, with accumulating evidence suggesting a role for cholesterol metabolism in HSC proliferation^53^. Mechanistically, our results indicate that PF4 limits the uptake of fluorescently labeled LDL by HSCs in a dose-dependent manner (**Figure 4E; supplemental Figure 4B**), suggesting an important role for PF4-LDLR in HSC metabolism. Next, we investigated if the expression of PF4 receptors was altered in old HSCs, which could contribute to the aging phenotype. Notably, aging did not affect CXCR3 and LDLR expression on HSCs, as seen by RNA and flow cytometry analysis (**supplemental Figures 4A and 4C**). As expected, old HSCs showed significantly increased uptake of LDL *in vitro* (**supplemental Figure 4D**), consistent with the notion that cholesterol accumulation is associated with the skewing of hematopoiesis toward myeloid lineages, a phenomenon observed in HSC aging^54,55^. Importantly, PF4 inhibited LDL uptake by old HSCs in a dose-dependent manner (**Figure 4F; supplemental Figures 4E**).

Next, we further characterized the *in vivo* hematopoietic compartments of *Ldlr^−/−^* and *Cxcr3^−/−^* mice. Since HSCs from young *Pf4^-/-^* mice show accelerated aging phenotypes (**Figure 2; supplemental Figure 2**), we hypothesized that deleting PF4 receptors would lead to a similar *in vivo* phenotype. Yet, we did not observe any significant change in the frequency and absolute number of HSCs in the bone marrow of *Ldlr^−/−^* or *Cxcr3^−/−^* mice, as seen in *Pf4^-/-^* mice (**supplemental Figures 4F-4G**) or myeloid-lineage bias (**supplemental Figure 4H**). To bypass a potential compensatory role between these two receptors, we crossed *Ldlr^−/−^* and *Cxcr3^−/−^* mice to generate double knockout mice (DKO mice; **Figure 4G**). DKO mice display smaller body sizes (data not shown) than WT, *Ldlr^−/−^* or *Cxcr3^−/−^* mice. Analysis of the hematopoietic compartment revealed that 3-month-old DKO mice present increased myeloid bias and reduced B cells in the bone marrow compared to control WT mice (**Figure 4H**). Interestingly, there is a marked increase in the frequency and number of phenotypic HSCs (**Figures 4I-4J; supplemental Figure 4I-4J**), alongside an increase in myeloid-primed progenitors in the bone marrow of DKO mice (**Figure 4K**). Accordingly, colony-forming assays revealed an increase in CFU-C activity, with a significant increase in granulocyte, macrophage, erythroid and megakaryocyte (GEMM) and granulocyte and macrophage (GM) colonies (**Figure 4L; supplemental Figure 4K**). Notably, HSCs from DKO mice displayed accelerated markers of HSC aging, as seen in *Pf4^-/-^* mice, including elevated levels of DNA damage identified by γH2AX staining (**Figure 4M**). Accordingly, cdc42 and a-tubulin polarity were significantly reduced (**Figures 4N** and **4O**), suggesting reduced regeneration and an aging-like phenotype ^37–41^. Overall, the similarities between HSC aging phenotypes in *Pf4^-/-^* mice and DKO mice, alongside the lack of response of *Ldlr^−/−^* and *Cxcr3^−/−^* HSCs to the PF4 signal, suggest that PF4 signals through LDLR and CXCR3 in HSCs.

### PF4 regulates the proliferation and lineage output of human HSCs

The age-related decline of HSCs occurs in both mice and humans^5–7^. Clinically, HSCs from older donors are associated with worse non-relapse mortality, highlighting the importance of developing HSC rejuvenation strategies^56^. Given our findings that PF4 can reverse aging phenotypes in murine HSCs, we next investigated whether similar effects could be achieved in human HSCs. To determine the optimal concentration of human recombinant PF4, we cultured human CD34+ cells under *ex vivo* conditions (**Figure 5A**) previously described^57^. Human phenotypic Lin^-^ CD34^+^ CD38^-^ CD90^+^ CD45RA^-^ HSCs exhibited a dose-dependent response to PF4, with 2.5 μg/mL being sufficient to induce maximum suppression of proliferation (**Figure 5B**). Cell cycle analysis further confirmed that PF4 induces quiescence in human HSCs (**Figure 5C**), consistent with observations in mice^11^. Indeed, PF4 could inhibit the expansion of human HSCs from different donor ages (ranging from 21 to 55 years old; **Figures 5D-5E; supplemental Figures 5A-5B**). To investigate the human HSC rejuvenation potential of PF4 further, we evaluated the HSC polarity and DNA damage, two age-related hallmarks of HSC aging^6^, in human middle-aged/old HSCs. Strikingly, PF4 treatment reduced DNA damage (**Figure 5F**) and increased the frequency of polarized HSCs (**Figures 5G-5H**). These findings suggest that PF4 also restores youthful characteristics in middle-aged/old human HSCs.

**Figure 5.**
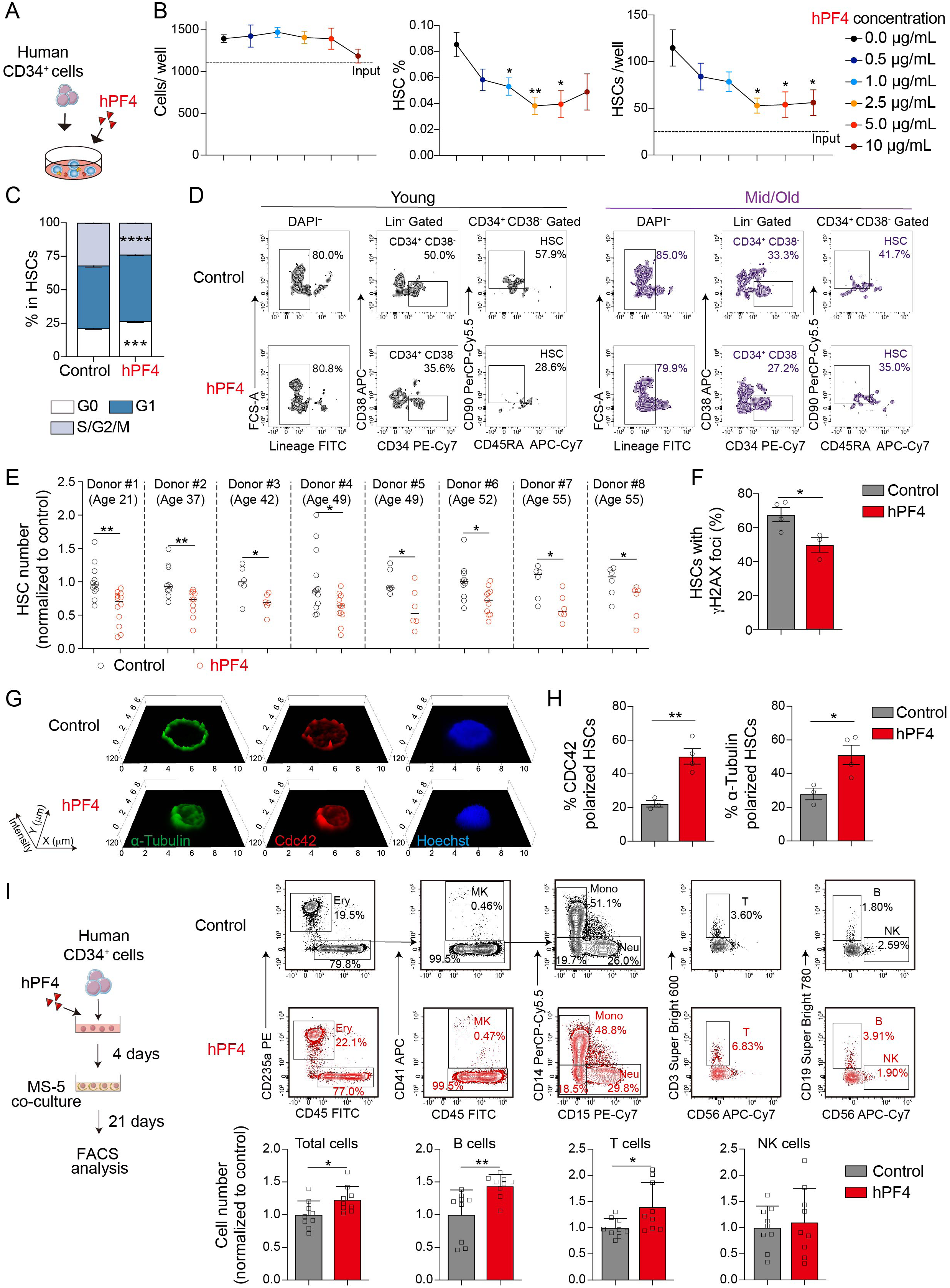
hPF4 regulates the proliferation of human HSCs. **A.** Schematic of human CD34^+^ BM cells cultured with recombinant human PF4 (hPF4). **B.** Number of human cells per well after cultured with hPF4 at the different concentrations (n = 6 wells for each condition). **C.** Cell cycle analysis of human HSCs treated with hPF4 (n = 6 wells for each condition). **D.** Representative FACS plots of human HSCs after hPF4 culture. **E.** Number of phenotypic HSCs (Lin^-^ CD34^+^ CD38^-^ CD90^+^ CD45RA^-^) per well, normalized to the control, after culture with hPF4 at 2.5 μg/mL (n = 6 - 12 wells for each donor). **F.** Percentage of HSCs (ages: 52-55) with γH2AX foci after culture with hPF4 for 18 hours (total of 80 and 52 cells from control and hPF4 treatment group). **G.** Representative confocal 2.5 D images of HSCs from the control and hPF4 treatment group stained with cdc42, α-Tubulin, and Hoechst. **H.** Quantification of the percentage of cdc42 and α-Tubulin polarized HSCs out of total HSCs scored (total of 86 and 97 cells from control and hPF4 treatment group). **I.** Experimental outline of the co-culture system (left graph). Representative FACS plots showing the differentiation of human HSCs (ages 52 and 55) into various cell types (top graph). Normalized cell numbers of total, B, T, and NK cells (n = 9, derived from 2 independent assays, bottom graph). Data are shown as mean ± SD. *P < 0.05, **P < 0.01, ***P < 0.001, ****P < 0.0001.

For functional validation, we performed a long-term culture assay with middle-aged/old human CD34^+^ cells following four days of PF4 treatment. The cells were co-cultured with the MS-5 cell line; a condition permissive to human multilineage hematopoietic differentiation^57,58^. After 21 days, flow cytometry analysis revealed that PF4 treatment significantly enhanced hematopoietic output, particularly lymphoid differentiation, with increased CD3^+^ T cells and CD19^+^ B cells (**Figure 5I**). Myeloid lineage output remained unchanged (**supplemental Figure 5C**). These results demonstrate that human HSCs are responsive to PF4 signaling, and PF4 treatment can restore lymphoid output in aging HSCs, suggesting the conservation of the PF4-mediated rejuvenation signal across species.

## Discussion

In recent years, considerable progress has been made in understanding the biology of MK ontogeny^59^ and how they further contribute to hematopoietic homeostasis during early development^60^ and adulthood by directly regulating HSC function^11,12,15,61,62^. However, the interactions between HSCs and MKs in the bone marrow niche are lost during aging^17–19^. Here, we demonstrate that age-related decline in the megakaryocytic niche capacity, marked by an immature phenotype, heightened pro-inflammatory signature, and decreased secretion of PF4, is an important driver of HSC aging. These changes can result in HSC exhaustion, diminished regenerative capacity, and a shift toward myeloid-biased differentiation, as MKs are the main niche regulators of the HSC myeloid-biased subset^15^. Our findings suggest that targeting MK-driven pathways, especially PF4, may offer a promising strategy to mitigate HSC aging and improve hematopoietic function in the elderly.

Several cell intrinsic^16^ and extrinsic mechanisms, such as attrition of the MK-associated vascular beds^63,64^, inflammation^17,28–31^, and alterations in adrenergic signals^17,18^ of the aged bone marrow, might be driving the age-related changes to megakaryopoiesis. Nevertheless, our results demonstrate that old MKs or the loss of MK-derived PF4 directly drives HSC aging, and administration of recombinant PF4 can partially reverse molecular and functional aging phenotypes.

Targeting distinct cell-intrinsic and extrinsic mechanisms has been proposed to rejuvenate aged HSCs with different levels of success^8^. While monotherapy might not be sufficient to fully rejuvenate an old hematopoietic system, the potential synergist effect of HSC-intrinsic and niche-based rejuvenation strategies holds great promise. Remarkably, recently, it was revealed that the administration of systemic PF4 could restore age-associated cognitive function. Though the precise PF4 cognitive rejuvenation mechanisms are still not fully understood, the mitigation of neuroinflammation is regarded as a possible explanation, suggesting a potential link between hematopoietic and neural aging^36,65,66^. Our study revealed that PF4 could restore aged HSCs to a balanced lineage output, benefiting immune cell function. Furthermore, systemic PF4 administration could improve HSC function, ultimately enhancing the brain’s regenerative capacity^67^. These findings highlight the importance of exploring HSC rejuvenation strategies, which will also exert neuroprotective effects.

Recently, it was shown that eliminating aged myeloid-biased HSCs via antibody-mediated depletion restored features of a more youthful immune system by enabling the activity of balanced HSCs and indirectly by reducing the production of inflammatory myeloid cells^68^. Mechanistically, it is likely that PF4, by suppressing the proliferation of myeloid-biased HSCs^15^, allows the activity of lineage-balanced HSCs to repopulate the hematopoietic system.

The receptors CXCR3 and LDLR play important roles in mediating the effects of PF4 on various cell types. While PF4 is essential in regulating HSCs quiescence and reconstitution potential, the absence of a single receptor (CXCR3 or LDLR) does not fully recapitulate the phenotype observed in PF4-deficient mice, potentially highlighting a compensatory role. CXCR3-deficient mice show a slight increase in HSC numbers with reduced self-renewal ability, while LDLR-deficient mice exhibit minimal alterations in hematopoiesis^69,70^. Interestingly, we found that double-knockout LDLR and CXCR3 could replicate the accelerated aging phenotype of PF4-deficient mice, suggesting that these receptors are critical for transmitting the PF4 signal in HSCs. Mechanistically, our results indicate that PF4 limits HSC proliferation by inhibiting LDL uptake, a mechanism associated with myeloid skewing^54,55^, and a phenomenon observed in HSC aging. CXCR3 activation can trigger multiple signaling pathways in distinct cell types^71^ however, the PF4-CXCR3 downstream signaling pathway in HSCs remains to be defined.

Our study demonstrates that phenotypic human HSCs also respond to the PF4 youthful signal. Yet, the expression levels of LDLR and CXCR3 are very low in human HSCs (data not shown), suggesting the interesting possibility that PF4 might act on distinct human receptors. Notably, PF4 treatment enforced HSC quiescence on HSCs isolated from different age groups, restored lymphoid lineage output, improved cell polarity, and decreased the frequency of HSCs with γH2AX foci. This preclinical evidence supports the potential for developing therapeutic approaches to rejuvenate aged donor HSCs via PF4 administration. However, further preclinical and clinical research will be essential to assess the feasibility and effectiveness of this therapy in humans. These findings could provide valuable insights into enhancing the health and function of older individuals by preventing or mitigating age-associated immune dysfunction and hematopoietic diseases.

## Acknowledgments

The authors thank Professor Mortimer Poncz for sharing the *Pf4^-/-^* mice and the UIC Flow Cytometry core facility for assistance with FACS sorting and the UIC Fluorescence Imaging Core for assistance with confocal imaging. The authors recognize and thank funding support for their laboratories from NIH (R01HL162584 to S.P.), (R01HL170286 to C.C.), (R01HL174801 to M.M.), a Longevity Impetus Grant from the Norn Group (to S.P. and C.C.), the American Society of Hematology (ASH) Scholar Award (to M.M.), and a Glenn Foundation for Medical Research Postdoctoral Fellowship in Aging Research (to S.Z.). A.M.D. was partially supported by an NIH predoctoral (F31HL165859) and an NIDDK TL1 Chicago KUH Forward training (TL1DK132769) fellowships. C.E.A. and C.M.P. were partially supported by an NIH training grant (T32HL007829).

## Authorship Contributions

S.Z. designed the study, performed most experiments, and analyzed the data. C.E.A., A.M.D., and C.M.P. helped with experiments. K.J. and E.H.W.L. performed and analyzed RNA-seq experiments. S.G.O. advised on the experimental setup and data interpretation. C.N. provided the *vWF-eGFP* mice. M.M. helped with the imaging experiments and advised on the experimental setup and data interpretation. S.P. and C.C. designed, supervised, and obtained funding for the study. S.Z., C.C., and S.P. wrote the manuscript. All authors discussed the results and commented on the manuscript.

